# Same-Species Contamination Detection with Variant Calling Information from Next Generation Sequencing

**DOI:** 10.1101/531558

**Authors:** Tao Jiang, Martin Buchkovich, Alison Motsinger-Reif

## Abstract

**Motivation:** Same-species contamination detection is an important quality control step in genetic data analysis. Compared with widely discussed cross-species contamination, same-species contamination is more challenging to detect, and there is a scarcity of methods to detect and correct for this quality control issue. Same-species contamination may be due to contamination by lab technicians or samples from other contributors. Here, we introduce a novel machine learning algorithm to detect same species contamination in next generation sequence data using support vector machines. Our approach uniquely detects such contamination using variant calling information stored in the variant call format (VCF) files (either DNA or RNA), and importantly can differentiate between same species contamination and mixtures of tumor and normal cells.

**Methods:** In the first stage of our approach, a change-point detection method is used to identify copy number variations or copy number aberrations (CNVs or CNAs) for filtering prior to testing for contamination. Next, single nucleotide polymorphism (SNP) data is used to test for same species contamination using a support vector machine model. Based on the assumption that alternative allele frequencies in next generation sequencing follow the beta-binomial distribution, the deviation parameter ρ is estimated by maximum likelihood method. All features of a radial basis function (RBF) kernel support vector machine (SVM) are generated using either publicly available or private training data. Lastly, the generated SVM is applied in the test data to detect contamination. If training data is not available, a default RBF kernel SVM model is used.

**Results:** We demonstrate the potential of our approach using simulation experiments, creating datasets with varying levels of contamination. The datasets combine, in silico, exome sequencing data of DNA from two lymphoblastoid cell lines (NA12878 and NA10855). We generated VCF files using variants identified in these data, and then evaluated the power and false positive rate of our approach to detect same species contamination. Our simulation experiments show that our method can detect levels of contamination as low as 5% with reasonable false positive rates. Results in real data have sensitivity above 99.99% and specificity at 90.24%, even in the presence of DNA degradation that has similar features to contaminated samples. Additionally, the approach can identify the difference between mixture of tumor-normal cells and contamination. We provide an R software implementation of our approach using the defcon()function in the vanquish: Variant Quality Investigation Helper R package on CRAN.

## 1. Introduction

High throughout next generation sequencing (NGS), has shown its advantages over traditional Sanger sequencing and microarrays in terms of accuracy, cost and speed (Metzker, 2010; van Dijk, Auger, Jaszczyszyn, & Thermes, 2014). As NGS technologies have matured, best practices for quality control and data processing procedures have also developed (Patel & Jain, 2012). One key step in a robust pipeline is the detection of sample contamination detection. Sample contamination detection is a necessary data quality control step of NGS data analysis pipeline since contamination might be introduced during sample preparation and sequencing analysis. Sample contamination affects all downstream sample analysis and may even generate misleading results, which in turn lead to false positive associations and genotype misclassification (Jun et al., 2012).

In this paper, our working definition of contamination is when any sample contains tissues from more than one contributing source. There are two major types of contamination: (1) cross-species contamination; (2) same-species or within-species contamination. Additionally, it is very common for a single cancer sample to contain both normal and tumor cells from the same patient or cell line.

Contamination can emerge in next generation sequencing samples for various reasons. Despite best practices, unclean lab devices are one of the most common causes, with the introduction of unexpected materials like *Mycoplasma* (Schmidt, Hummel, & Herrmann, 1995). This same problem exists in even more recent, large scale research projects, such as the 1000 Genomes Project (Langdon, 2014). Contamination can also arise from sample handling, sample extraction, library preparation and amplification, sample multiplexing and inaccurate barcode sequencing (Simion et al., 2018). Available methods are mainly based on the sequencing and allele frequency information of the testing sample and can be categorized into two groups, according to the source of contaminants: cross-species contamination and same-species contamination.

Cross-species contamination has been well studied, with several methods that have emerged to detect this type of contamination (Korneliussen, Albrechtsen, & Nielsen, 2014; Laurence, Hatzis, & Brash, 2014; Schmieder & Edwards, 2011; Strong et al., 2014). In fact, modern metagenomics approaches are extensions of cross species contamination detection approaches. For example, Schmieder and Edwards developed DeconSeq, a framework for identification and removal of human contamination from microbial metagenomes (Schmieder & Edwards, 2011), providing a framework for cross-species contamination detection during sequencing alignment. Later in 2014, Merchant *et al.* used microbiome analysis software to scan a sample of domestic cow, *Bos taurus,* and found small contigs from microbial contaminants (Merchant, Wood, & Salzberg, 2014). Generally, data is assembled from available Sanger reads to the known species, then the unmapped contigs within the assembly are classified by *k*-mer matching where a database containing all bacteria, archaea, and viruses from the RefSeq database. As a result, contigs aligning to other genomes are found as a sign of contamination.

By contrast, same-species or within-species contamination is relatively more challenging, with fewer methods to implement valid and robust approaches. Arguably the most commonly implemented approach, and the earliest developed is ContEst (Cibulskis et al., 2011), a module within the Genome Analysis ToolKit (GATK) software (McKenna et al., 2010). ContEst uses a Bayesian method to calculate the posterior probability of each contamination level and find the maximum *a posterior* probability (MAP) estimate of the contamination level at homozygous loci. Assuming a uniform prior distribution, Unif(0,1), on the contamination level, the posterior distribution of contamination level is proportional to the joint distribution of seeing observed alleles given the qualities of base calling and the probabilities of observing true alleles in contaminated sample. Thus, ContEst requires VCF and BAM format input and general population frequency information, i.e., base identities and quality scores from sequencing data.

Later in 2012, Jun et al. developed the VerifyBamID package for detecting same-species contamination of human DNA samples in both sequence and array-based data (Jun et al., 2012). VerifyBamID implements both likelihood-based and regression-based approaches that assume no more than one contaminant contained in a testing DNA sample. The likelihood of a contamination level is maximized by using a grid search over each contamination level for a maximizer. While VerifyBamID has demonstrated good sensitivity in real normal data experiments, copy number alterations (CNAs) in tumor samples will shift allele frequencies away from those outside of CNA regions, which leads to misinterpreting copy number-driven shift as contamination (Bergmann, Chen, Arora, Vacic, & Zody, 2016a).

The statistical model introduced in VerifyBamID, was further developed by Bergmann *et al.* in their Conpair method for detecting another source of same-species contamination - the mixture of tumor and normal cells from the same patient sample (Bergmann, Chen, Arora, Vacic, & Zody, 2016b). Conpair focuses on homozygous loci within tumor-normal paired samples. Given that homozygous markers are invariant to copy number changes; pre-selected highly informative genomic homozygous markers are provided with Conpair to perform contamination detection.

Other more recent methods developments have used haplotype structure for contamination detection in NGS data (Sehn et al., 2015). Closely spaced SNP pairs within sequencing region are identified from the 1000 Genomes database (The 1000 Genomes Project Consortium, 2012). Then read haplotypes for these selected SNP pairs are inferred. Human-human admixture is suggested if more than two read haplotypes are observed at a given locus in a sample. The estimated level of contamination for each testing sample are twice the mean frequency of the minor haplotype.

While the current approaches have been successful in a broad range of applications, there are major limitations that we address with our current approach. These developments represent substantial improvements in both the practical implementation of the quality control procedures and the statistical model used. Current approaches rely on the sizeable human reference genome data, as well as at least two large, memory intensive files. They each require either tumor and normal BAM files (Conpair), or VCF file and BAM file (VerifyBamID and ContEst). By using a combination of a Beta-Binomial assumption and support vector machines within our algorithm to detect same species contamination, it works directly from information in VCF file. Even for a tumor-normal paired sample, no other information is needed to perform contamination detection. The change points of B allele frequencies (from the VCF file) are detected and then all chromosomes are separated into shorter sequences. Each sequence overlapping any copy number variation or abberation region is detected and filtered. In the enclosed study, we demonstrate our method in both real and simulated data, and show that it demonstrates excellent sensitivity and specificity for both simulated and real data. We also describe an R package implementation of the method.

## 2. Materials and Methods

### 2.1. Beta-binomial Model of Allele Frequency in Next Generation Sequencing

The proposed method is designed for human applications and assumes a diploid genome. For each locus that contains a single nucleotide variant (SNV) called from next generation sequencing (NGS) data, we define the allele frequency as the number of counts for the alternative (non-reference genome) allele over the total number of depth. For any diploid genome, if an individual is homozygous for the alternative allele (denoted as alternative/alternative, 1/1), the expected allele frequency is 1; meanwhile, if an individual is heterozygous (denoted as reference/alternative, 0/1) at a locus, 0.5 is the expected allele frequency. Using these theoretical expectations motivates using the Binomial distribution for the number of reads at each locus,

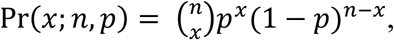

where *n* is the total number of depth at the locus; *p* is the theoretical allele frequency; *x* is the number of counts for the alternative allele.

While such a simple model is intuitively *appealing*, previous studies have discovered extra binomial dispersion, specifically, overdispersion of allele frequency distributions (Esteve-Codina et al., 2011; Pickrell et al., 2010; Skelly, Johansson, Madeoy, Wakefield, & Akey, 2011; Zhang et al., 2014). This overdispersion results in a higher variability than the Binomial distribution, so a distribution that models such large variance is needed. Previous studies proposed and demonstrated the Beta-Binomial distribution as an appropriate model for allele frequencies at a particular locus in a subpopulation. The Beta-binomial distribution is a discrete hierarchical model containing the Beta distribution and binomial distribution, where the probability follows the Beta distribution; and the response follows the Binomial distribution. Hence the probability mass function of the Beta-Binomial distribution is

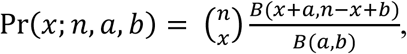

where *n* is the total number of reads at the locus; *B(a, b)* is the beta function theoretical allele frequency; *x* is the number of counts for the alternative allele. This model has been applied in a number of studies since, and the advantages of the Beta-Binomial over the Binomial distribution in dealing with overdispersion has been repeatedly demonstrated (Chen et al., 2016; Mayba et al., 2014). This work motivates our use of the Beta-Binomial distribution.

### 2.2. Quality Control of Variant Call Format (VCF) files

The input format for our method is the well-established Variant Call Format (VCF< Danecek et al., 2011). To our knowledge, this is the first method to detect same-species contamination from the VCF format. The VCF format contains all single nucleotide variation (SNV) information that is needed in the next steps. Before a VCF file is used in our software, quality control is needed to filter noise and unnecessary information. The recommended quality control and processing steps are outlined below, and are additional quality control steps beyond the processing to produce the VCF file itself.

#### Step 1. Indel Filtering

The SNVs used in our method must be substitution variants, not copy number variations like insertions and deletions. Only substitution mutations result in heterozygous and homozygous genotypes that are appropriately modeled by the Beta-Binomial distribution. Insertions and deletions are identified as any mutation segments with a length of more than one base pair, and are then filtered/dropped in this step.

#### Step 2. Homozygous and Heterozygous Genotype Calling

The next step of processing is to call the genotypes for modeling. As mentioned above, there are two genotypes for alternative allele at any SNV: homozygous and heterozygous. Suggested genotypes are listed in the GT field of VCF file where 0/0 means homozygous reference; 0/1 means heterozygous; and 1/1 means homozygous alternative. In step 2 of processing, new information is generated that summarizes the genotype in reference to the alternative allele, in which “Het” reflects 0/1 in GT, and “Hom” reflects 1/1 in GT. This results in two categories of called variants, each corresponding to its own Beta-binomial model. Homozygous reference (0/0) and other heterozygous genotypes (1/2, 2/3, and so on) are not considered to simplify computations, and are label as “Complex” and are not included in further calculations.

#### Step 3. Low and High Depth Filtering

NGS data is not completely error free, so it is important to identify whether a sequence is a true call or a sequencing error. One of the efficient methods is to set thresholds for coverage depth (Morgan et al., 2010). A previous study suggested read depths of >50 provide acceptable sensitivity and specificity in mutation detection. A reasonable read depth threshold should be chosen according to the average read depths of a testing sample.

#### Step 4. Change point Detection for CNV

In a pure sample, when there is a copy number variation (CNV) region, its features look very similar to the region with more than one contributors (same species contamination). Hence it is necessary to filter CNV region before generating features. If there is CNV information already generated for testing sample, the function vanquish::defcon() can filter the CNV region directly. Otherwise, a change point detection method is used to detect the CNV region. It has been reported that the variances of B-Allele Frequency (BAF, alternative allele frequency) at heterozygous loci are different among normal, duplication, deletion and loss of heterozygous (Ku et al., 2013). Therefore, change point analysis can be employed to detect change point of variance, i.e., border of copy number region. The package “changepoint” is applied on only heterozygous positions for multiple changepoint searching of variance change (Rebecca Killick & Eckley, 2014).

### 2.3. Distribution and Likelihood based Features

The next step of our approach is to generate variables/features that will be used for model building to predict whether a sample contains same-species contamination. Briefly, there are two types of features that are generated and used in model building: distribution-based feature and likelihood-based feature.

#### (1)Distribution based features

By its definition, allele frequency is a real number between 0 and 1. To generate distribution-based features, allele frequency is binned into 4 regions as shown in **Figure 1**: Low Alternative Allele Frequency (LowRate), Heterozygous Alternative Allele Frequency (HetRate), High Alternative Allele Frequency (HighRate), and Homozygous Alternative Allele Frequency (HomRate). Here 0, 0.3, 0.7, and 0.99 are their corresponding cut-off values.

**Figure 1.**
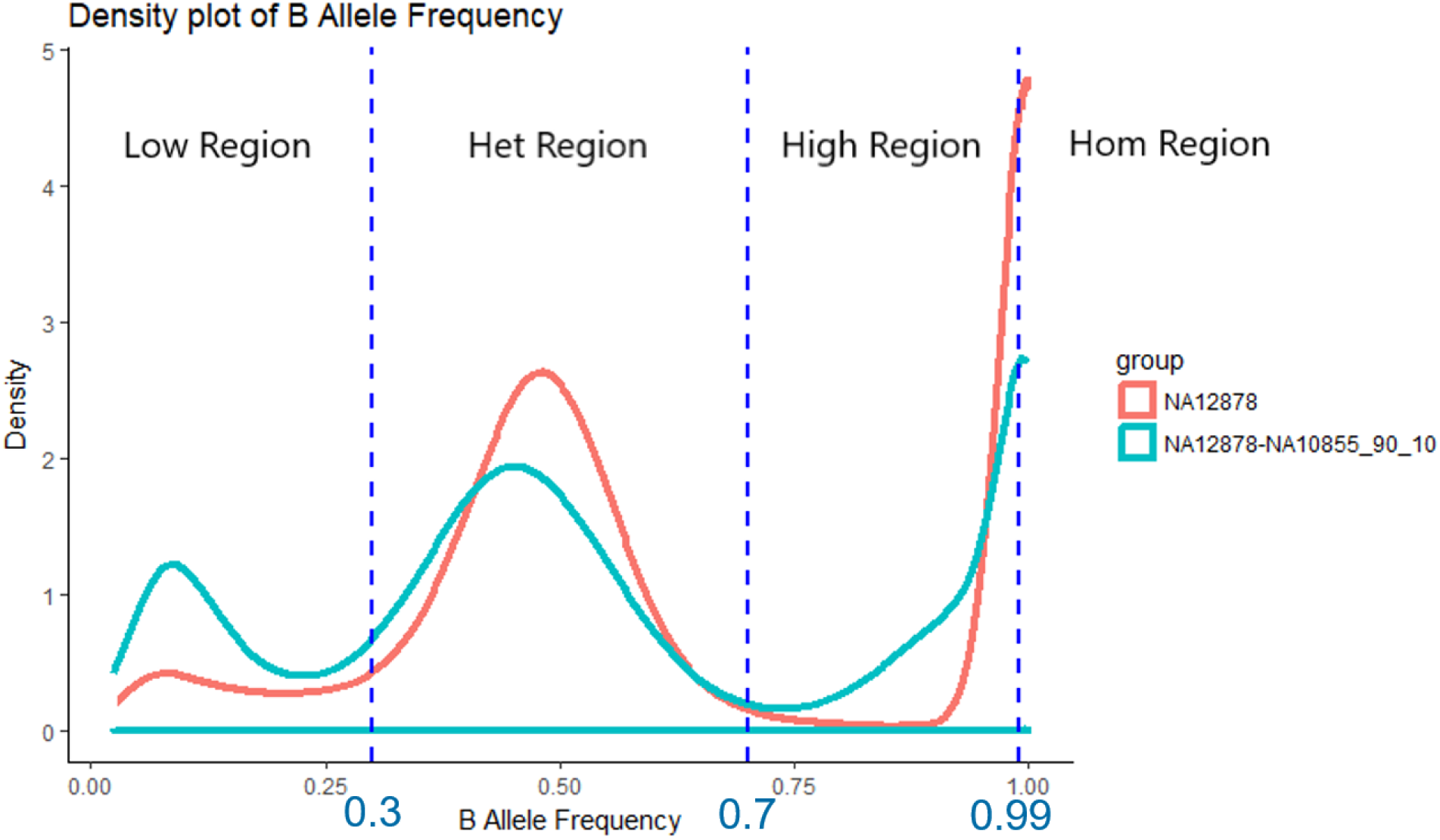
Four regions for B-Allele frequency: LowRate, HetRate, HighRate, and HomRate and their corresponding cut-off values. LowRate is the region where B-Allele frequency is below 0.3; HetRate is the region where B-Allele frequency is between 0.3 and 0.7; HighRate is the region where B-Allele frequency is between 0.7 and 0.99; HomRate is the region where B-Allele frequency is above 0.99. **B Allele frequency density plots of pure (NA12878) and contaminated (NA12878-NA10855_90_10) samples.** The features used in SVM classifier are based on the difference between pure and contaminated curves.

From the bins in **Table 1**, there are 8 distribution-based features generated for the following model-building steps. These features reflect the distribution of allele frequencies in the whole file, instead of at each variant calling position. Therefore, each input sample/ VCF file has one set of features to represent itself.

**Table 1.**
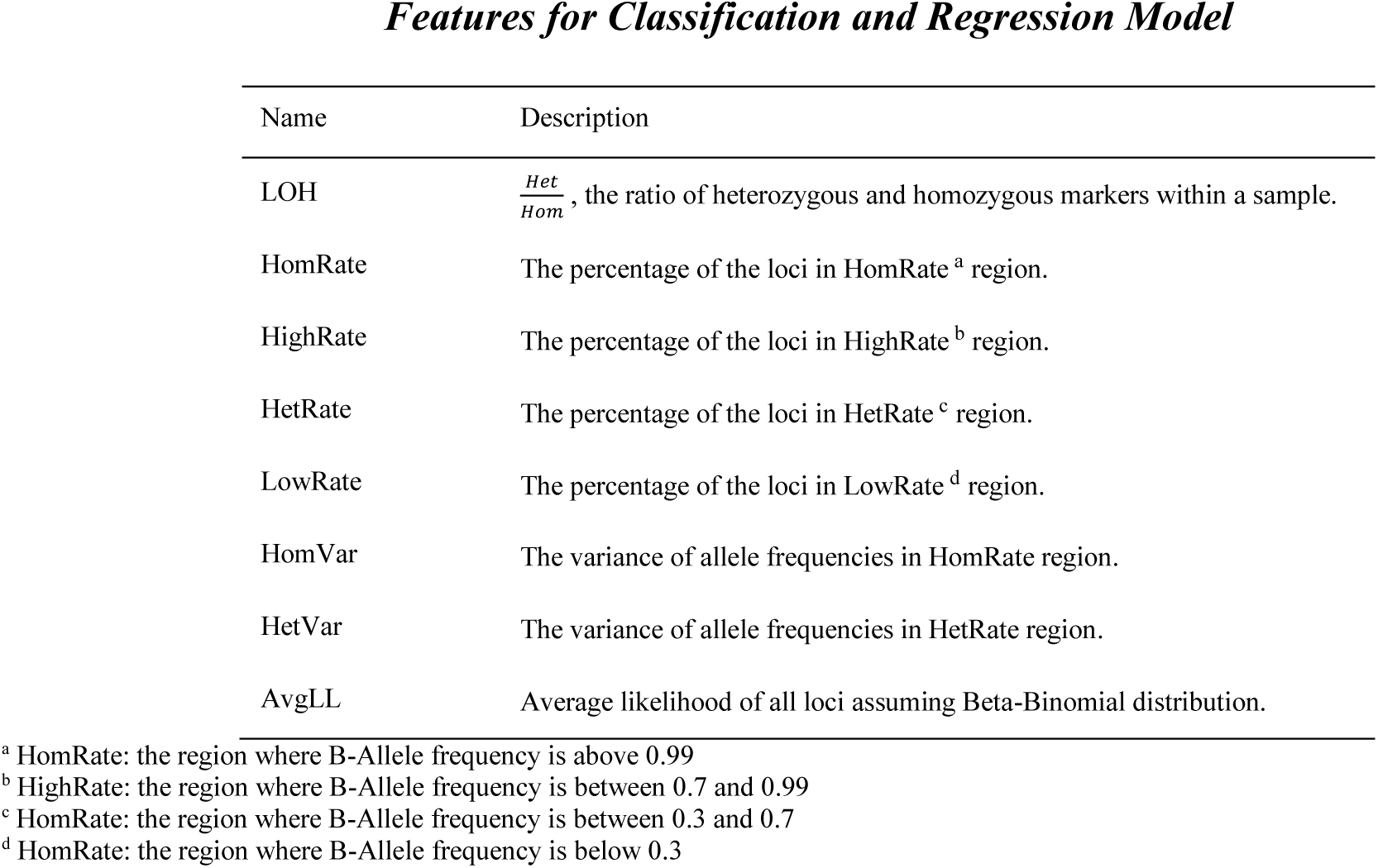
All features for classification and regression model and their descriptions.

*Features for Classification and Regression Model*
| Name | Description |
| --- | --- |
| LOH | $\frac{Het}{Hom}$, the ratio of heterozygous and homozygous markers within a sample. |
| HomRate | The percentage of the loci in HomRate <sup>a</sup> region. |
| HighRate | The percentage of the loci in HighRate <sup>b</sup> region. |
| HetRate | The percentage of the loci in HetRate <sup>c</sup> region. |
| LowRate | The percentage of the loci in LowRate <sup>d</sup> region. |
| HomVar | The variance of allele frequencies in HomRate region. |
| HetVar | The variance of allele frequencies in HetRate region. |
| AvgLL | Average likelihood of all loci assuming Beta-Binomial distribution. |

<sup>a</sup> HomRate: the region where B-Allele frequency is above 0.99
<sup>b</sup> HighRate: the region where B-Allele frequency is between 0.7 and 0.99
<sup>c</sup> HomRate: the region where B-Allele frequency is between 0.3 and 0.7
<sup>d</sup> HomRate: the region where B-Allele frequency is below 0.3

#### (2)Average log-likelihood, the likelihood based feature

The only likelihood based feature is the average likelihood of all loci in VCF file. The likelihood is calculated by applying the Beta-binomial distribution. In the current study, NA 10855 and other available pure samples are selected as a reference genome to calculate maximum likelihood estimator for both parameters, *p* and *ρ*, in the Beta-binomial distribution (Sequenced at Q2 Solutions). With 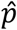 and 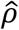 log likelihood of all loci are calculated so that the last feature, their average value is also generated.

### 2.4. Support Vector Machine

After generating features, a classification method is used to identify whether a sample is from single contributor or multiple contributors. In the current study, we apply a Support Vector Machine (SVM) model because of the complexity in pattern recognition within feature space (Cortes & Vapnik, 1995). The SVM method fits a hyperplane between single and multiple contributor regions to optimally discriminate between classifications. In our approach, we implement the SVM method using the e1071 R package (Meyer et al., 2018) (https://CRAN.R-project.org/package=e1071). Since a linear model is not guaranteed, the Gaussian (Radial Basis Function) kernel is used to avoid such parametric assumptions. There are two parameters in SVM analysis that need to be tuned: cost and gamma. They are tuned using the parallel searching method, where a grid search is conducted on an exponentially growing sequence of cost and gamma to look for optimized paired values. It is possible for the estimated parameter to be different, given another training data set.

## 3. Results

### 3.1. Changepoint analysis for approximate copy number region detection

In the scenario where the copy number information of a sample is not provided, change point analysis was conducted to find copy number regions of a sample. The rmChangePoint() function within the vanquish package imports cpt.var() from changepoint package (Rebecca Killick & Eckley, 2014). The pruned exact linear time (PELT) method (R. Killick, Fearnhead, & Eckley, 2012) and the Changepoints for a Range of PenaltieS (CROPS) algorithm (Haynes, Eckley, & Fearnhead, 2014) are employed to search for variance changepoints. In **Figure 2(A)**, the B-allele frequencies (only those between 0.05 and 0.95) of corresponding loci contained within the input VCF files are plotted. It is clear that there exist some CNV patterns. Red vertical lines are where the variance changes detected. The whole plot is separated into multiple parts by these change points. For each part, if the percentage of loci of the B-allele frequency is between 0.45 and 0.55 is larger than 10% and the skewness is larger than 0.5, the part will be kept in further analysis. The result after filtering is shown in **Figure 2(B)**. See the documentation of the vanquish package for more details.

**Figure 2.**
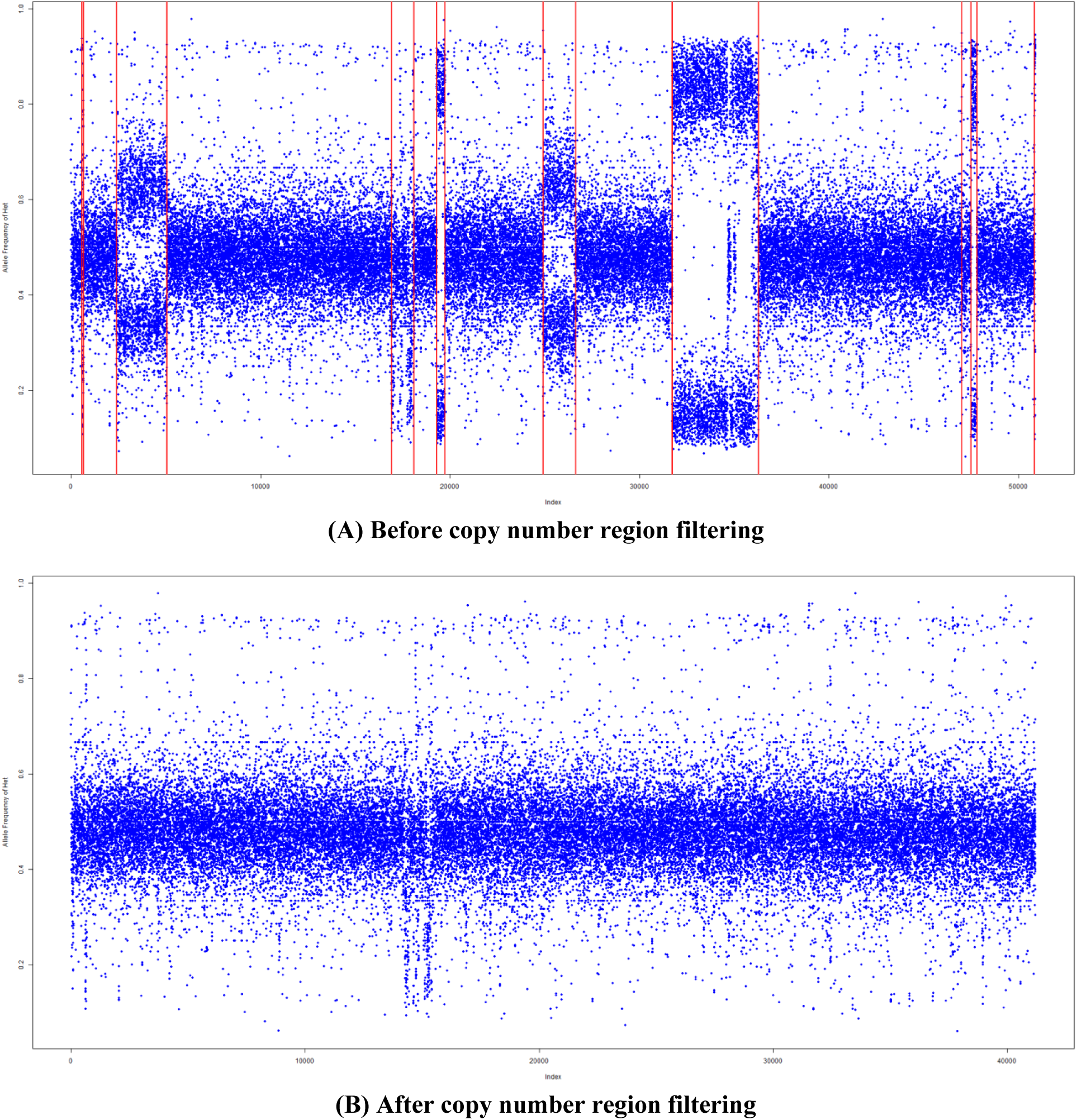
Change point analysis for copy number region detection. The Y-axis shows B-Allele frequency; X-axis is shows the location number of each variant from chromosome 1 to 22. (A) is before copy number region filtering. Red lines indicate where the variance change based on detection results. It is clear that there exist some copy number patterns. (B) is after copy number region filtering and in which most copy number patterns are removed.

### 3.2. Beta-binomial parameter estimation for reference sample(s)

In order to calculate likelihood-based features for further analysis, maximum likelihood estimators of *ρ*for beta-binomial distribution of heterozygous and homozygous models are estimated. For the B-allele frequency, the theoretical value of the parameter pis 0.5 in heterozygous model and 1 in homozygous model. pis fixed at 0.5 and 0.999 to search for*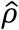* in corresponding model. L-BFGS-B (Byrd, Lu, Nocedal, & Zhu, 1995) is applied for maximum value searching. For instance, NA10855 was chosen as a reference sample, and five replicates were sequenced by Q2 Solutions. The maximum likelihood estimator of *ρ* in each sample was estimated by rho_est() in the vanquish package (**Table 2**) and the sample averages were achieved for further analysis. The value the of estimator highly depends on the variant caller, so it is suggested to keep using the same variant caller for the reference sample, training sample, and test samples.

**Table 2.** Maximum Likelihood Estimator 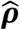 of NA10855 Samples. 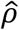 of heterozygous and homozygous models were estimated for each sequencing replicate. The sample mean can be used for generating features from training data set.

| | Heterozygous $\hat{\rho}$ | Homozygous $\hat{\rho}$ |
| --- | --- | --- |
| NA10855_1 | 0.154 | 0.0269 |
| NA10855_2 | 0.223 | 0.0253 |
| NA10855_3 | 0.177 | 0.0210 |
| NA10855_4 | 0.187 | 0.0310 |
| NA10855_5 | 0.169 | 0.0274 |
| **Sample Mean** | **0.182** | **0.0263** |

### 3.3. Features in the classification and regression model

To train the classification and regression model, 238 samples were sequenced at Q2 Solutions as a training data set. 124 out of the 238 samples were pure; and the rest of the training set were contaminated. Some of the contaminated samples were mixed on purpose in wet lab; and others were simulated by two pure FASTQ format files (Cock, Fields, Goto, Heuer, & Rice, 2010). Pure and contaminated samples show different B-allele frequency patterns. As shown in **Figure 3**, only heterozygous loci detected in samples are plotted. The pure sample (**Figure 3A**) had a narrow horizontal band; while the contaminated sample (**Figure 3B**) had a relatively uniform distribution for B-allele frequency. Eight boxplots and t-tests (null hypothesis ofno differences) were conducted to show the difference of pure and contaminated samples considering each feature **(Figure 4)**. Among all eight features after t-tests, HomVar, HetVar and HighRate have significant p-values 1.786^-9^,, 1.750^-6^, and 4.540^-20^ correspondingly. See Supplementary data for details of each feature.

**Figure 3.**
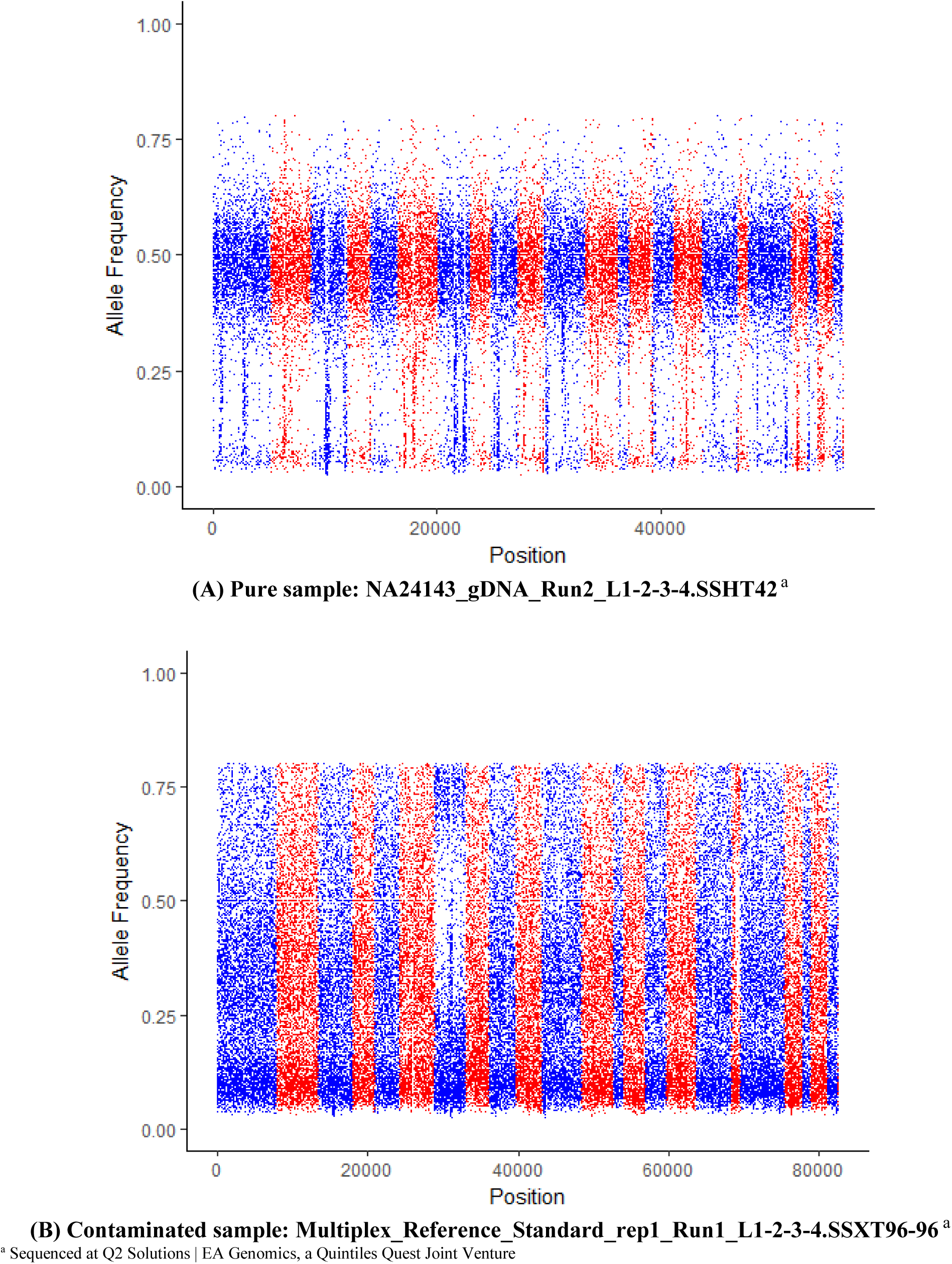
Heterozygous B-allele frequency patterns of pure (A) and contaminated (B) samples. Blue and red bands indicate different chromosomes.

**Figure 4.**
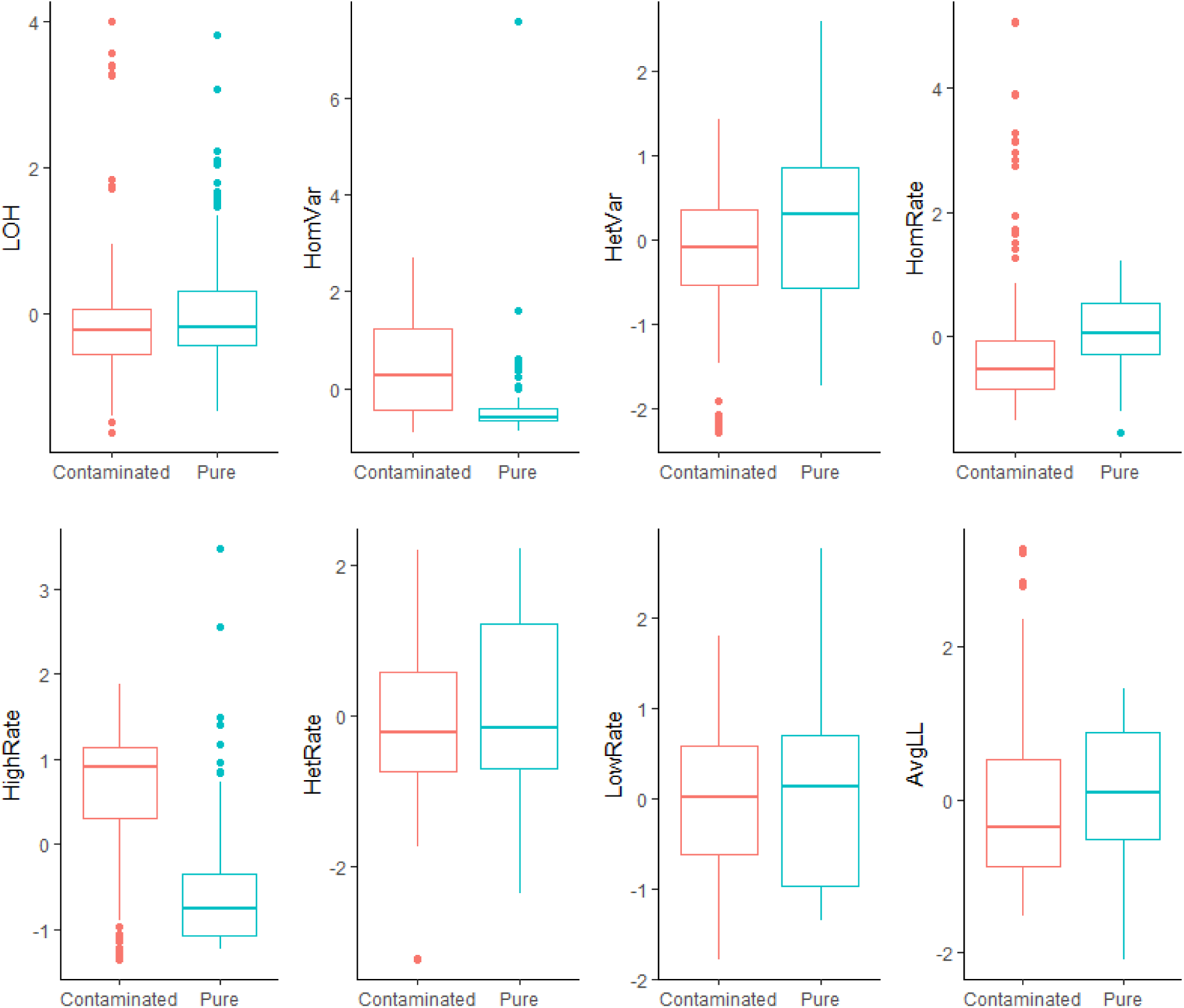
Boxplots of all features for pure samples (124) vs contaminated samples (114). Among all eight t-tests (null hypothesis of no difference hypothesis), HomVar, HetVar and HighRate have significant p-values 1.786^-9^,, 1.750^-6^, and 4.540^-20^ correspondingly.

### 3.4. Tune cost and gamma parameters in radial kernel SVM

Monte Carlo method (1000 times) and tune() from R package e1071 (Meyer et al., 2018) (https://CRAN.R-project.org/package=e1071) were used to tune parameters cost and gamma in support vector machine. All 238 samples were split into training (70%, 167 samples) and test (30%, 71 samples) sets. For the training set, we used grid search to tune parameter cost in the range of (2^-4^, 2^12^), and gamma in the range of (2^-4^, 2^4^). Then sensitivity and specificity were calculated from the test set by using tuned cost and gamma. The results of Monte Carlo simulation are in **Table 3**. Median values of cost (16) and gamma (0.25), mean values of sensitivity (97.65%) and specificity (96.27%) were presented and the tuned cost and gamma were used in radial kernel SVM model for prediction.

**Table 3.**
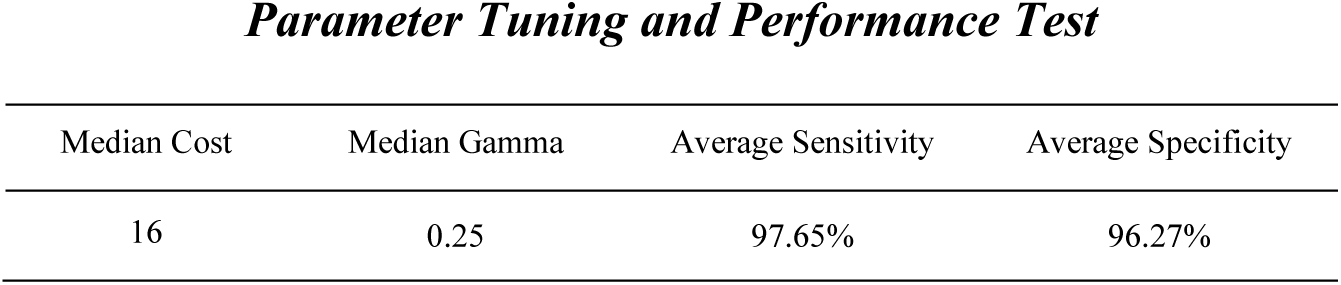
Monte Carlo test results for parameter tuning and performance test. *Contamination Detection for Simulated Data*

| Median Cost | Median Gamma | Average Sensitivity | Average Specificity |
| --- | --- | --- | --- |
| 16 | 0.25 | 97.65% | 96.27% |

### 3.5. Simulated data test results

Two reference samples, NA12878 and NA10855, were sequenced at Q2 Solutions. Two pairs of FASTQ format files were obtained from sequencing results, and where re-sampled and mixed according to different proportions as shown in **Table 4** by seqtk (https://github.com/lh3/seqtk). In this simulated test, NA12878 was treated as the sample, while NA10855 was assumed to be mixed into the NA12878 sample. The percentages of the artificial contaminated mixtures range from 0.5% to 20%, and the detection rate at different levels of contamination were calculated. The total number of reads were ∼50 million for all 6 mixture samples. After running contamination analysis, contamination percentages above 5% were readily detected, while lower percentages were not detectable (**Table 4**). Based on this result, the contamination detection analysis has a detection sensitivity down to 5% in this case. If contaminant is less similar to the sample that it is mixed with, the detection sensitivity will be lower; on the other hand, if the similarity is larger, then contamination detection analysis will be more challenging.

**Table 4.** Contamination Detection for a series of simulated data. *Contamination Detection for Real Data*

| Sample Component | Reads (NA12878) | Reads (NA10855) | Test Results |
| --- | --- | --- | --- |
| NA12878 (80%) + NA10855 (20%) | 40M <sup>a</sup> | 10M | Contaminated |
| NA12878 (90%) + NA10855 (10%) | 45M | 5M | Contaminated |
| NA12878 (95%) + NA10855 (5%) | 47.5M | 2.5M | Contaminated |
| NA12878 (97.5%) + NA10855 (2.5%) | 48.75M | 1.25M | Pure |
| NA12878 (99%) + NA10855 (1%) | 49.5M | 0.5M | Pure |
| NA12878 (99.5%) + NA10855 (0.5%) | 49.75M | 0.25M | Pure |

<sup>a</sup> M: million

### 3.6. Two groups of real data test results

After quantitative simulation test, the trained model was applied on a real data set (22 samples in all); and the results are shown in **Table 5**. The samples are ranked by regression values from e1071::svm(). Twenty out of twenty-two samples had the same prediction as prior identification. However, a pair of Human_T-Lymphoblast samples (see bold text in **Table 5**) were considered as contaminated but predicted as pure. Therefore, we zoomed in to check the distribution of B-allele frequency in this pair of samples (**Figure 5**). The copy number variation pattern (two bands and three bands in chromosomes) was shifted below (middle area of CNV pattern shifted from 0.5 to 0.3) since the sample was a mixture of tumor normal cells from the same person. The pattern was clear: the source of sample was uniform. Based on the distance of shift, the percentage of tumor cells and normal cells of the sample is even predictable.

**Table 5.** Contamination Detection for a series of real data. Twenty out of twenty-two samples had the same prediction as prior identification. However, a pair of Human_T-Lymphoblast samples (bold text) were considered as contaminated but predicted as pure.

| Sample Name | Classification | Regression | Prior Identification |
| --- | --- | --- | --- |
| Human_B-Lymphocyte_L8 | 1 | 1.9243094 | 1 |
| Human_B-Lymphocyte_2_L20 | 1 | 1.9209875 | 1 |
| Human_Breast_2_L16 | 0 | 1.483925 | 0 |
| Human_Breast_L4 | 0 | 1.463376 | 0 |
| **Human_T-Lymphoblast_2_L21<sup>a</sup>** | **0** | **1.3622305** | **1** |
| **Human_T-Lymphoblast_L9** | **0** | **1.3472358** | **1** |
| Human_Brain_L3 | 0 | 1.3147938 | 0 |
| Human_Brain_2_L15 | 0 | 1.303287 | 0 |
| Human_Testis_L12 | 0 | 1.245767 | 0 |
| Human_Cervix_2_L17 | 0 | 1.2429423 | 0 |
| Human_Testis_2_L24 | 0 | 1.2424441 | 0 |
| Human_Cervix_L5 | 0 | 1.203943 | 0 |
| Human_Macrophage_L10 | 0 | 1.158416 | 0 |
| Human_Macrophage_2_L22 | 0 | 1.1582528 | 0 |
| Human_Liver_2_L18 | 0 | 1.1442246 | 0 |
| Human_Liposarcoma_L7 | 0 | 1.1406007 | 0 |
| Human_Liposarcoma_2_L19 | 0 | 1.132044 | 0 |
| Human_Skin_2_L23 | 0 | 1.1209464 | 0 |
| Human_Skin_L11 | 0 | 1.1194772 | 0 |
| Human_Liver_L6 | 0 | 1.1170909 | 0 |
| Human_Reference_DNA_Male_L1 | 0 | 1.0945151 | 0 |
| Human_Reference_DNA_Male_2_L13 | 0 | 1.0906951 | 0 |

<sup>a</sup> This is a mixture of tumor and normal cells. See **Figure 5** for distribution of B-allele frequency of this sample.

**Figure 5.**
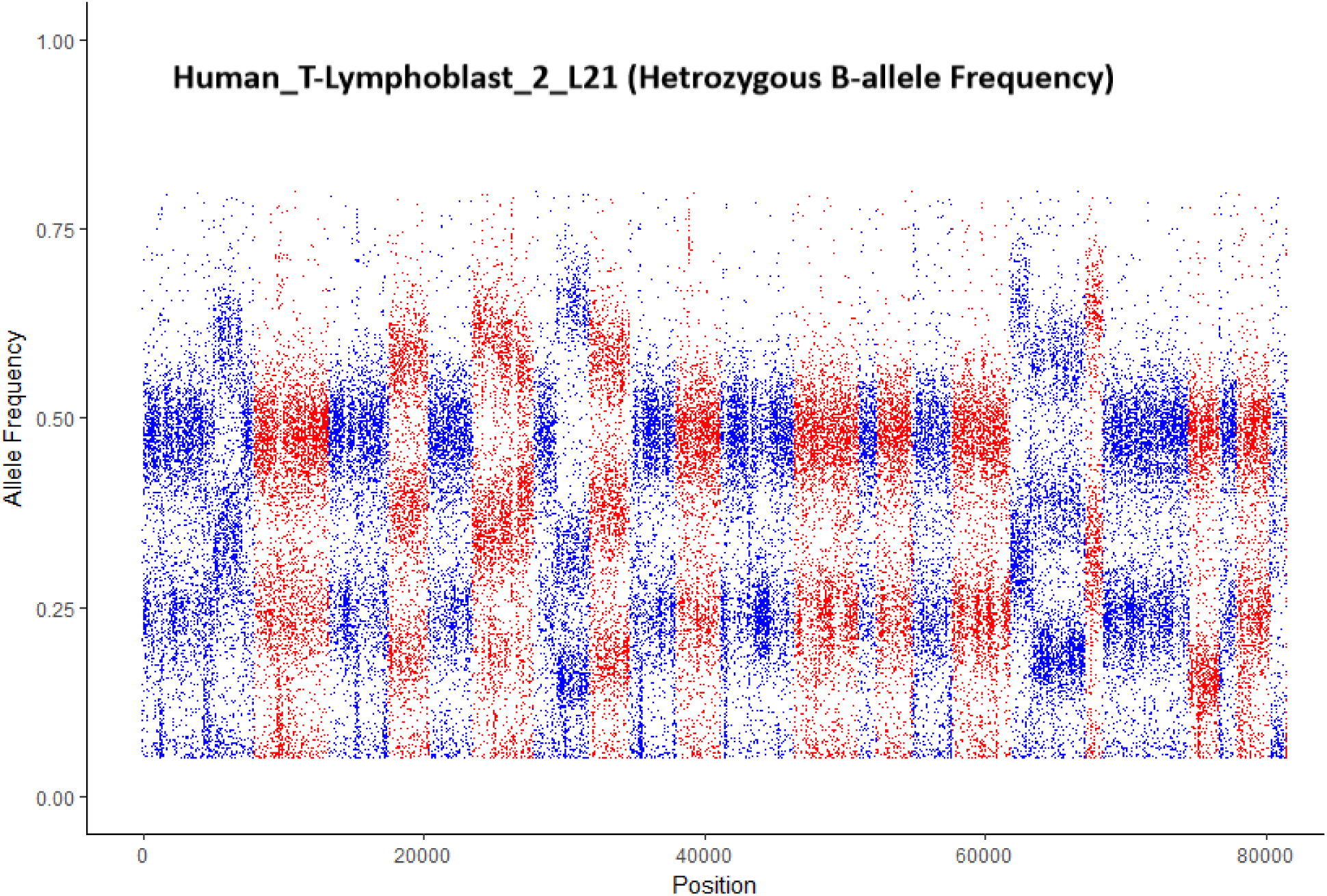
A sample that was previously considered as contaminated, but was detected as pure by vanquish::defcon(). The copy number variation pattern was shifted lower (middle area of CNV pattern shifted from 0.5 to 0.3) since the sample was a mixture of tumor normal cells from the same person.

A second data set (contains 53 samples in all) was also used to test the model (See **supplementary information** for more details). Within this data set, 12 samples were mixed with contaminant on purpose; 41 samples were pure. The test results show that the sensitivity is > 99.99%, and the specificity is 90.24%. All four false positive samples were formalin-fixed paraffin-embedded (FFPE) tissues, and were very likely to be degraded (**Figure 6**). When generating features from this degraded sample in **Figure 6**, its features looked very similar to features from a contaminated sample.

**Figure 6.**
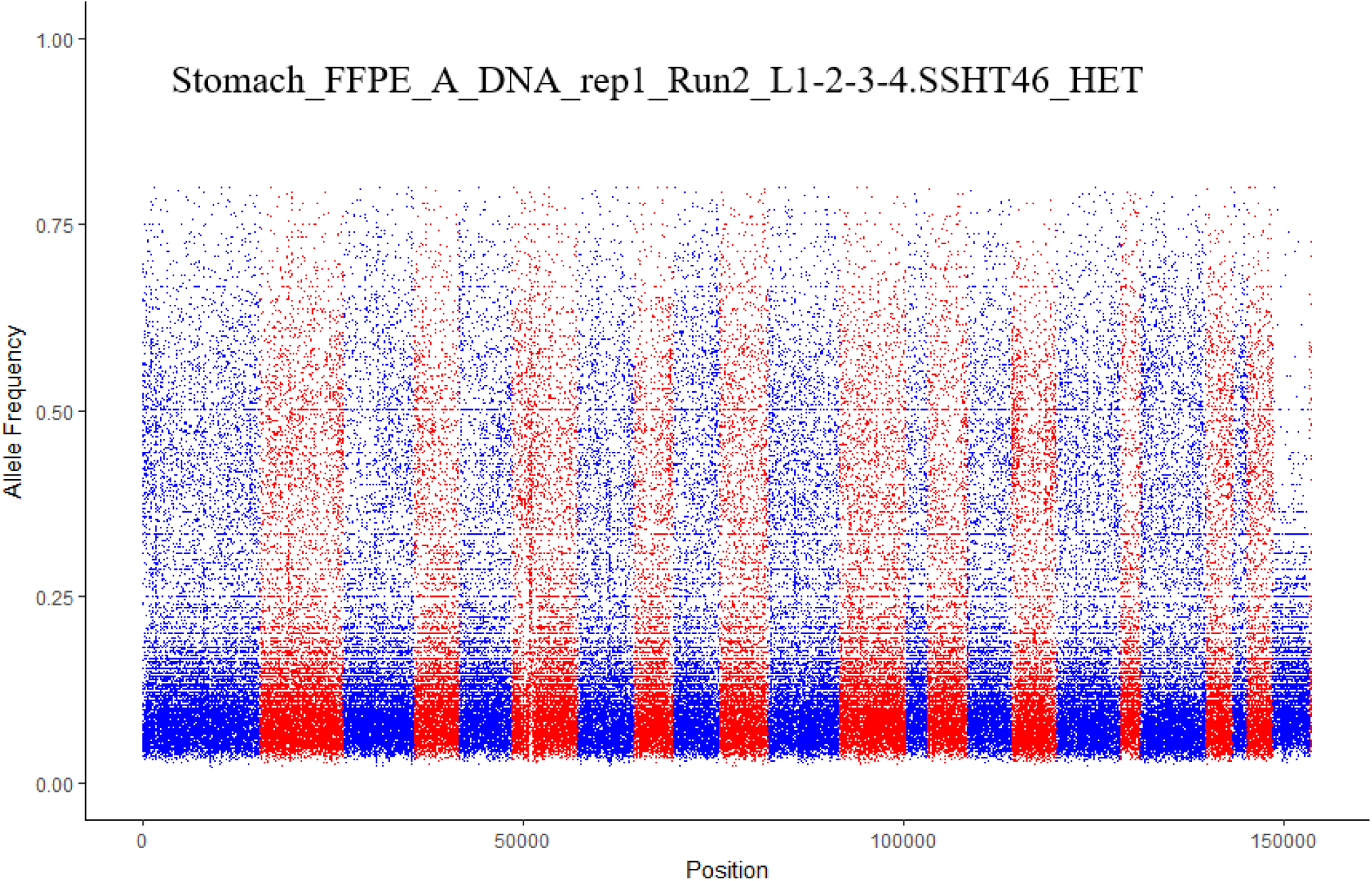
Formalin-fixed paraffin-embedded (FFPE) tissue with low quality. This sample was a false positive prediction result because of the similarity between its low quality and contaminants.

## 4. Discussion

In this study, we introduced a novel strategy that detected same-species or within-species contamination by using B-allele frequency from variant call information only. An R package, vanquish: Variant Quality Investigation Helper, was produced to conduct this analysis. A brief flowchart (**Figure 7**) introducing contamination detection procedure shows the necessary steps as following:

**Figure 7.**
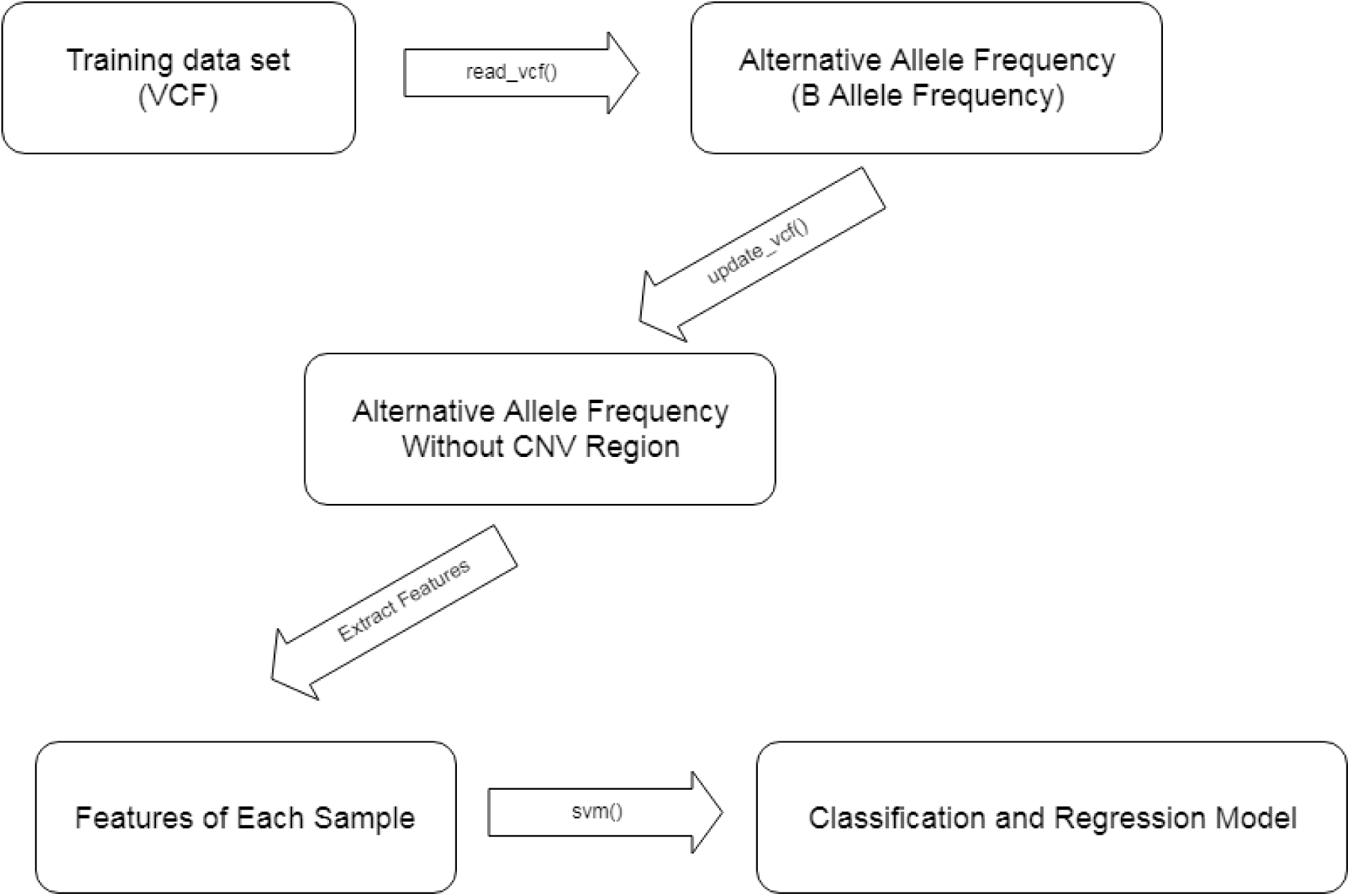
A brief flowchart of contamination detection procedure.

1. Variant call format (VCF) generated by a variant caller was read into R by vanquish::read_vcf function. So far, the supported variant callers are GATK, VarDict and strelka2.
2. Copy number variation regions existing in VCF file were approximately detected and filtered by vanquish::update_vcf function.
3. Features for radial kernel SVM model were extracted from each sample by vanquish::generate_feature function.
4. Parameter cost and gamma for kernel SVM were tuned.
5. A test sample was predicted whether it was contaminated.

Additionally, two scenarios were found where the performance of this approach may be affected. First, if the sample is a mixture of tumor and normal cells from the sample individual, which is not considered contamination by our definition. The second scenario is when the test sample has very low quality so that no clear B-allele frequency pattern can be found.

Finally, we have produced a user friendly R package to enable rapid analysis for same species analysis. Our tool uniquely performs this important step of QC from VCF files, improving the performance, memory requirements, etc. The running time of CNA region removal and feature generating is summarized in **Figure 8** (Hardware: Dell R820, 512GB of RAM). Five samples without any known change point are run 10 times each to achieve their average running time, keeping their maximum numbers of runs of algorithm are uniform. Same as our expectation, larger samples need more time to do change point detection and feature generation. Besides, if a sample contains more change points, it also needs a longer running time.

**Figure 8.**
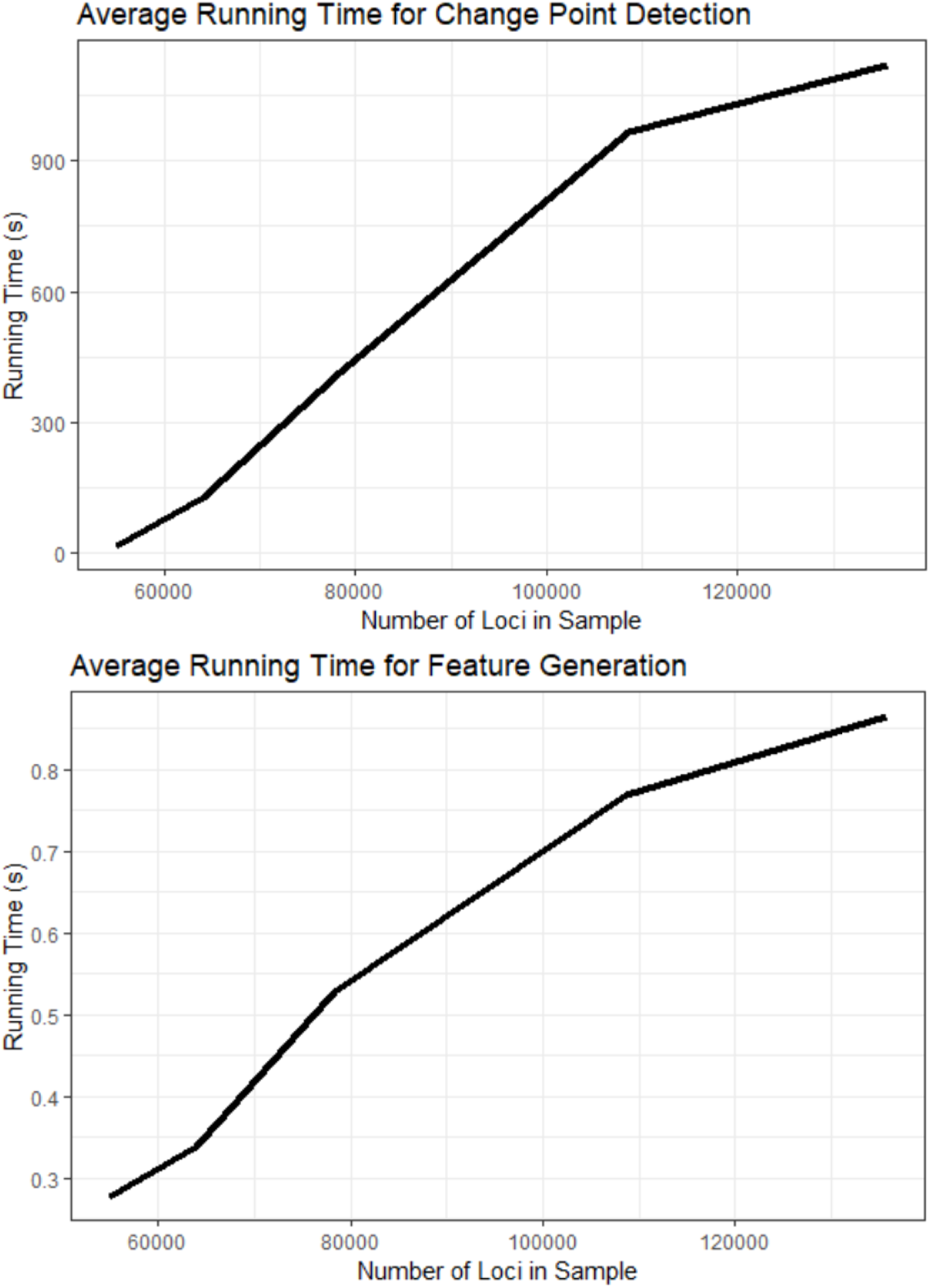
Average running time (of 10 runs) for change point detection (Top) and feature generation (Bottom). Five samples with different numbers of loci but the same maximum number of runs of algorithm (= 2).

## Acknowledgements and funding

This work was supported by Q2 Solutions | EA Genomics, a Quintiles Quest Joint Venture and P01 CA142538 from the National Cancer Institute. The authors also gratefully thank Dr. Chad Brown for discussions and inspiration on this contamination detection project.

