## Supplemental Information for "Same-Species Contamination Detection with Variant Calling Information from Next Generation Sequencing"

SUPPLEMENTARY METHODS

### Features in Support Vector Machine (SVM)

*LOH: Loss of Homozygous*

All SNPs distributed across chromosomes are classified either homozygous (1/1) or heterozygous (0/1). LOH is the ratio of the number of heterozygous SNP loci over the number of homozygous SNP loci. A large LOH value means that there are more heterozygous SNP loci in a sample.

*HomeRate / HighRate / HetRate / LowRate*

Each SNP locus has a respective B-Allele frequency (BAF), which is percentage of the depth alternative allele out of the total depth at each SNP locus. Here BAF $\in [0,1]$, and three cut-off values have been applied to separate the support set of BAF, [0,1], into 4 sub-regions: HomRate region $[0.99, 1]$, HighRate region $[0.7, 0.99)$, HetRate region $[0.3, 0.7)$, and LowRate region $[0, 0.3)$. A pure sample is expected to have higher HomeRate and HetRate values than a contaminated sample.

HomRate is the number of loci with BAF $\in [0.99, 1]$ over the total number of SNP loci in a sample.

HighRate is the number of loci with BAF $\in [0.7, 0.99)$ over the total number of SNP loci in a sample.

HetRate is the number of loci with BAF $\in [0.3, 0.7)$ over the total number of SNP loci in a sample.

LowRate is the number of loci with BAF $\in [0, 0.3)$ over the total number of SNP loci in a sample.

*HomVar / HetVar*

For SNP loci distributed within the HomRate region as defined above, the variance of BAF values of these SNP loci is calculated and defined as HomVar. HetVar is calculated in a similar procedure. A pure sample is expected to have lower HomeVar and HetVar values than a contaminated sample.

*AvgLL: Average Log-likelihood*

BAF of a SNP locus is assumed to follows the Beta-Binomial distribution. A reference sample (NA10855 sequenced at Q2 Solutions) is assumed as a pure sample. The chosen reference sample is used to calculate the maximum likelihood estimators for parameters $p$ and $\rho$ in the Beta-Binomial distribution. After that, log-likelihood of all SNP loci in testing sample are summed up. For comparability purpose, the sum of log-likelihood is then divided by the number of loci in each sample so that the final outcome is the average value of log-likelihood across all loci in a sample. A pure sample is expected to have higher AvgLL value than a contaminated sample.

### The Radial Basis Function Kernel in Support Vector Machine (SVM)

Let $\{(\mathbf{x}_1, y_1), \ldots, (\mathbf{x}_N, y_N)\}$ be a training data set formed by $N$ VCF files (training samples), where $\mathbf{x}_i$ is a vector of $d$ features ($d$ = 8 here) extracted from a VCF file, $y_i = \{+1, -1\}$ is the marked contamination label of the $i$th sample ($+1$ for contaminated sample; and $-1$ for pure sample), and $N$ is the size of training data set. As the definition of SVM, a linear discriminant function of an SVM is

$$f(\mathbf{x}) = \mathbf{w}^{\mathrm{T}}\mathbf{x} + b,$$

where $\mathbf{x}$ is from a training sample, $\mathbf{w}$ is the weight vector, and $b$ is the bias, both are parameters to be estimated. This can be formulated as optimizing using hinge loss as the loss function:

$$\min_{\mathbf{w}\in\mathbb{R}^{\boldsymbol{d}}} \|\mathbf{w}\|^2 + C\sum_{i}^{N} \max_{\mathbf{w},b}(0,1 - y_i f(\mathbf{x}_{\boldsymbol{i}}))$$

over $\mathbf{w}$ and $b$, where $C$ is the soft margin constant, a regularization parameter. For the purpose of transformation to kernel space, we focus on dual SVM classifier. By Representer Theorem, the solution of $\mathbf{w}$ can be written as a linear combination of the feature vectors and contamination labels, such as

$$\mathbf{w} = \textstyle\sum_{i}^{N} \alpha_i y_i \mathbf{x}_i.$$

Therefore, directly plugging $\mathbf{w}$ gives us a new classifier

$$\begin{aligned} f(\mathbf{x}) &= \left(\sum_{i}^{N} \alpha_i y_i \mathbf{x}_i\right)^{\mathrm{T}} \mathbf{x} + b \\ &= \sum_{i}^{N} \alpha_i y_i \, \mathbf{x}_i{}^{\mathrm{T}}\mathbf{x} + b \end{aligned}$$

where $\alpha_i$ is the weight of the $i$th training sample. Only those samples selected as support vector machines will have non-zero $\alpha_i$'s. Meanwhile, the new objective function to optimize over $\boldsymbol{\alpha} \in \mathbb{R}^{\boldsymbol{N}}$, for $0 \leq \alpha_i \leq C$, $1 \leq i \leq N$, $\sum_{i}^{N} \alpha_i y_i = 0$, is

$$\max_{\alpha_i} \textstyle\sum_{i}^{N} \alpha_i - \frac{1}{2}\sum_{j,k}^{N} \alpha_j \alpha_k y_j y_k (\mathbf{x}_j{}^{\mathrm{T}}\mathbf{x}_k).$$

In a transformed feature space, define $\Phi(\mathbf{x})$ as a feature map, such that $\Phi\colon \mathbf{x} \longrightarrow \Phi(\mathbf{x}), \mathbb{R}^{\boldsymbol{d}} \longrightarrow \mathbb{R}^{\boldsymbol{D}}$, where $D$ is the dimension of $\Phi(\mathbf{x})$ in the transformed feature space. Hence the new discriminant function in the transformed feature space is

$$f(\mathbf{x}) = \sum_{i}^{N} \alpha_i y_i \, \Phi(\mathbf{x}_i)^{\mathrm{T}}\Phi(\mathbf{x}) + b$$

and the respective loss function is

$$\max_{\alpha_i} \sum_{i}^{N} \alpha_i - \frac{1}{2}\sum_{j,k}^{N} \alpha_j \alpha_k y_j y_k \Phi(\mathbf{x}_j)^{\mathrm{T}}\Phi(\mathbf{x}_k)$$

for $0 \leq \alpha_i \leq C$, $1 \leq i \leq N$, $\sum_{i}^{N} \alpha_i y_i = 0$.

Define the kernel function as $k(\mathbf{x}_j, \mathbf{x}_k) = \Phi(\mathbf{x}_j)^{\mathrm{T}}\Phi(\mathbf{x}_k)$, and so there is such a kernel classifier defined as

$$f(\mathbf{x}) = \sum_{i}^{N} \alpha_i y_i k(\mathbf{x}_i, \mathbf{x}) + b$$

and the respective loss function is

$$\max_{\alpha_i} \sum_{i}^{N} \alpha_i - \frac{1}{2} \sum_{j,k}^{N} \alpha_j \alpha_k y_j y_k k(\mathbf{x}_j, \mathbf{x}_k)$$

for $0 \leq \alpha_i \leq C$, $1 \leq i \leq N$, $\sum_i^N \alpha_i y_i = 0$.

Given the possible complexity of hyperplane for contamination detection problem, the Radial Basis Function (Gaussian) Kernel is chosen, for which the kernel function is

$$k(\mathbf{x}_j, \mathbf{x}_k) = \exp(-\gamma \|\mathbf{x}_j - \mathbf{x}_k\|^2),$$

where $\gamma > 0$ is the inverse-width parameter of Gaussian kernel. Therefore, the discriminant function and loss function are

$$f(\mathbf{x}) = \sum_{i}^{N} \alpha_i y_i \exp(-\gamma \|\mathbf{x}_j - \mathbf{x}_k\|^2) + b$$

$$\max_{\alpha_i} \sum_{i}^{N} \alpha_i - \frac{1}{2} \sum_{j,k}^{N} \alpha_j \alpha_k y_j y_k \exp(-\gamma \|\mathbf{x}_j - \mathbf{x}_k\|^2)$$

where $\gamma > 0$ , $0 \leq \alpha_i \leq C$, $1 \leq i \leq N$, $\sum_i^N \alpha_i y_i = 0$.

**Tunable hyper-parameters: *C* and $\gamma$**

Two hyper-parameters: the soft margin constant C and the inverse-width parameter of Gaussian kernel $\gamma$ are optimized by using grid search and cross validation. Grid search is applied on exploring the two dimensional space $(C, \gamma)$. The grid points of C are chosen on an exponential scale of $(2^{-4}, 2^{12})$; and grid points of $\gamma$ are chosen between $(2^{-4}, 2^{4})$. The sensitivity and specificity are estimated for each point on the grid.
